# Sclerostin modulates the degree of mineralization and the stiffness profile of the fibrocartilaginous enthesis for mechanical tissue integrity

**DOI:** 10.1101/2023.11.13.565114

**Authors:** Shinsei Yambe, Yuki Yoshimoto, Kazutaka Ikeda, Koichiro Maki, Aki Takimoto, Shinnosuke Higuchi, Xinyi Yu, Kenta Uchibe, Shigenori Miura, Hitomi Watanabe, Tetsushi Sakuma, Takashi Yamamoto, Kotaro Tanimoto, Gen Kondoh, Denitsa Docheva, Taiji Adachi, Chisa Shukunami

**Affiliations:** Department of Molecular Biology and Biochemistry, Division of Dental Sciences, Graduate School of Biomedical and Health Sciences, Hiroshima University, Hiroshima 734-8553, Japan; Department of Orthodontics and Craniofacial Developmental Biology, Applied Life Sciences, Graduate School of Biomedical and Health Sciences, Hiroshima University, Hiroshima 734-8553, Japan; Laboratory of Biomechanics, Institute for Life and Medical Sciences, Kyoto University, Kyoto 606-8507, Japan; Department of Maxillofacial Anatomy and Neuroscience, Division of Oral Health Sciences, Graduate School of Biomedical & Health Sciences, Hiroshima University, Hiroshima 734-8553, Japan; Laboratory of Integrative Biological Science, Institute for Life and Medical Sciences, Kyoto University, Kyoto 606-8507, Japan; Division of Integrated Sciences for Life, Graduate School of Integrated Sciences for Life, Hiroshima University, Higashi-Hiroshima, Hiroshima 739-8526, Japan; Department of Musculoskeletal Tissue Regeneration, Orthopaedic Hospital König-Ludwig-Haus, Julius-Maximilians-University Würzburg, 97074 Würzburg, Germany

**Keywords:** sclerostin, *Sost*, fibrocartilage, mineralization, fibrochondrocytes

## Abstract

Fibrocartilaginous entheses consist of four graded tissue layers including tendon, the unmineralized and mineralized fibrocartilage, and subchondral bone with varying degrees of stiffness. Here we examined the functional role of sclerostin that is expressed in mature mineralized fibrochondrocytes. Following rapid mineralization of the unmineralized fibrocartilage and parallel replacement of epiphyseal hyaline cartilage by bone, the unmineralized fibrocartilage re-expanded after a decline in alkaline phosphatase activity at the mineralization front. Sclerostin was co-expressed with osteocalcin in the bottom of the mineralized fibrocartilage adjacent to subchondral bone. In *Scx* deficient mice with less mechanical loading due to defects of the Achilles tendon, the number of sclerostin^+^ fibrochondrocytes was significantly reduced in the defective enthesis where chondrocyte maturation was markedly impaired in both fibrocartilage and hyaline cartilage. Loss of the *Sost* gene, coding for sclerostin, caused increased mineral density in the mineralized zones of the fibrocartilaginous enthesis. Atomic force microscopy analysis revealed the higher stiffness of fibrocartilage. These lines of evidence suggest that sclerostin in mature mineralized fibrochondrocytes acts as a modulator for mechanical tissue integrity of the fibrocartilaginous enthesis.

## INTRODUCTION

Cartilage is avascular, aneural, and alymphatic connective tissue that includes hyaline cartilage, elastic cartilage, and fibrocartilage (Shukunami et al., 2008; Takimoto et al., 2009; Hall, 2015). The most prevalent type is hyaline cartilage in which chondrocytes produce large amounts of type II collagen (Col2) and aggrecan (Acan). Hyaline cartilage includes the cartilaginous bone primordium that will serve as a template for the future bone during development as well as the articular cartilage that permanently covers to protect the epiphyseal bone surface at the joints (Hall, 2015). Fibrocartilage is found in the pubic symphysis, the inner annulus fibrosus of intervertebral discs, the knee menisci, the articular disc of the temporomandibular joint, and fibrocartilaginous entheses. Fibrocartilage has an appearance intermediate between hyaline cartilage and dense regular connective tissues such as tendons and ligaments (Benjamin and Evans, 1990; Hall, 2015).

The enthesis is the attachment site of the functional dense connective tissue components such as tendons, ligaments, joint capsule, or facia into bone in adult, but also to hyaline cartilage in fetal to childhood (Benjamin et al., 2006). Fibrocartilaginous entheses occur at the epiphysis or the apophysis of the bone, whereas fibrous enthesis directly attaches to the diaphysis (Sugimoto et al., 2013a; Apostolakos et al., 2014). We and others previously reported that primordial entheses arise from Scx^+^/Sox9^+^ progenitors during development to form as the junction between the dense connective tissue component and hyaline cartilage (Blitz et al., 2013; Sugimoto et al., 2013a).

In the fibrocartilaginous enthesis consisting of tendon, the unmineralized and mineralized fibrocartilage, and subchondral bone, the Gli1^+^ cell population is required for the mineralized fibrocartilage formation (Dyment et al., 2015; Schwartz et al., 2015). A recent single-cell RNA-seq analyses have unveiled distinct cell subpopulations in the fibrocartilaginous enthesis (Fang et al., 2022). Therefore, it is becoming increasingly important to verify specific cell populations at the protein level and to determine the spatiotemporal localization of each cell population within the tissue in in-depth studies.

The Achilles tendon, which is the strongest and largest tendon in the body, connects the soleus and gastrocnemius muscles to the calcaneus. At birth, the Achilles tendon, which is positive for tenomodulin (Tnmd), is attached to the calcaneus consisting of hyaline cartilage, which is positive for chondromodulin (Cnmd) (Sugimoto et al., 2013a). Fibrocartilaginous enthesis of the Achilles tendon develops postnatally in response to mechanical stimuli (Benjamin and Ralphs, 1998; Benjamin et al., 2006). Fibrochondrocytes rapidly mature to mineralize their surrounding matrix through the activation of hedgehog signaling (Dyment et al., 2015; Schwartz et al., 2015). Unlike mineralized hyaline cartilage except for the bottom zone of articular cartilage, mineralized fibrocartilage remains avascular and is not replaced by bone throughout life (Dyment et al., 2015). Once injured, the loss of a gradual mineral transition results in the decreased mechanical performance of the load-bearing interface, and healing process would not follow developmental processes that generate functionally graded layers of the fibrocartilaginous enthesis (Ideo et al., 2020). Although mechanical tissue integrity of the fibrocartilaginous enthesis would be coordinately regulated by combination of intrinsic factors and extrinsic mechanical forces transmitted via tendon, it is still uncertain how such a link is created during fibrocartilaginous enthesis formation.

Sclerostin, the *Sost* gene product, is a secreted protein that is predominantly expressed in osteocytes but also in articular hypertrophic chondrocytes (van Bezooijen et al., 2004; Poole et al., 2005; van Bezooijen et al., 2009). Sclerostin antagonizes canonical Wnt signaling and several BMP responses (Li et al., 2005; Krause et al., 2010). Sclerostin acts as a negative regulator of bone formation and favors bone resorption (Baron and Kneissel, 2013). *Sost* deficiency causes sclerostenosis and Van Buchem disease, autosomal recessive disorders (Brunkow et al., 2001; van Bezooijen et al., 2009). In this study, we analyzed the functional role of sclerostin in the postnatal development of the fibrocartilaginous enthesis of the Achilles tendon. For immunostaining and analysis by atomic force microscopy (AFM), we took advantage of the Kawamoto’s film method to use cryofilms for preparation of the thin fresh frozen sections from undecalcified hard tissues (Kawamoto, 2003; Takimoto et al., 2015; Kawamoto and Kawamoto, 2021). More antibodies can work on fresh, undecalcified sections without destroying the epitope by heat or organic solvents. Sclerostin expression was persistently detected in the mineralized mature fibrochondrocyte layer adjacent to subchondral bone. In *Scx* deficient mice with decreased mechanical loading due to defective tendon formation (Killian and Thomopoulos, 2016; Yoshimoto et al., 2017; Shukunami et al., 2018), both fibrocartilage and hyaline cartilage development were impaired and sclerostin expression was markedly decreased. Loss of *Sost* resulted in the increased bone mineral density of subchondral bone and the mineralized fibrocartilage. AFM analysis revealed that fibrocartilage stiffness is significantly higher in *Sost* deficient mice. Thus, sclerostin marking mature mineralized fibrochondrocytes modulates the degree of mineralization and the stiffness profile of the fibrocartilaginous enthesis for mechanical tissue integrity.

## RESULTS

### Expression of sclerostin in mature fibrochondrocytes of the mineralized fibrocartilage

In hyaline cartilage, Sox9^+^ chondrocytes produce Col2 and Acan and then mature to become hypertrophic chondrocytes synthesizing Col10 prior to mineralization (Kozhemyakina et al., 2015). Using Kawamoto’s film method for sectioning undecalcified hard tissues, we compared the expression of these cartilage markers in the calcaneus and its insertion site of the Achilles tendon by immunostaining (Fig. 1). Expansion of the unmineralized fibrocartilage and the underlying hyaline cartilage at P7 was visualized by toluidine blue (TB) staining (Fig. 1A). Col1 was co-expressed with Scx in tendon and fibrocartilage (Fig. 1B). Col2 was detected in both epiphyseal hyalin cartilage and fibrocartilage, while Scx was expressed in the upper portion of Col2^+^ fibrocartilage of the enthesis near tendon (Fig. 1C). Only a small number of alkaline phosphatase (ALP)^+^ cells were observed at the junction between the unmineralized fibrocartilage and epiphyseal hyaline cartilage (Fig. 1D). At P14, hypertrophic chondrocytes appeared in epiphyseal hyaline cartilage beneath fibrocartilage (Fig. 1E). ALP activity was high in osteoblasts, fibrochondrocytes and chondrocytes except for the resting hyaline cartilage (Fig. 1F) and chondroclasts/osteoclasts positive for tartrate-resistant acid phosphatase (TRAP) were found in the mineralized hypertrophic cartilage of the growth plate and the primary spongiosa (Fig. 1G). Intense Col2 staining was detected in Sox9^+^ cartilage and cartilage remnants around the chondro-osseus junction (Fig. 1H-J), whereas Col1 was co-expressed with Scx in fibrocartilage, tendon, and primary spongiosa (Fig. 1K). At P18, Sox9 was expressed in proliferating and resting chondrocytes (Fig. 1L,M). In hyaline cartilage, hypertrophic/mineralized chondrocytes strongly expressed Col10, whereas its expression in the mineralized fibrocartilage is faint (Fig. 1N,O). Thus, mineralizing chondrocytes are divided into two distinct groups: Col2^+^/Col10^+++^/Col1^-^ hypertrophic chondrocytes in epiphyseal hyaline cartilage and Col2^++^/Col10^+^/Col1^++^ fibrocondrocytes in the enthesis. We also found that sclerostin was expressed in mature fibrochondrocytes at P28 (Fig. 1P). Calcein labelling and ALP/Alizarin red (AR) staining revealed that the mineralization front consisting of ALP^+^ cells extends towards the midsubstance of the Achilles tendon (Fig. 1Q,R).

**Fig. 1.**
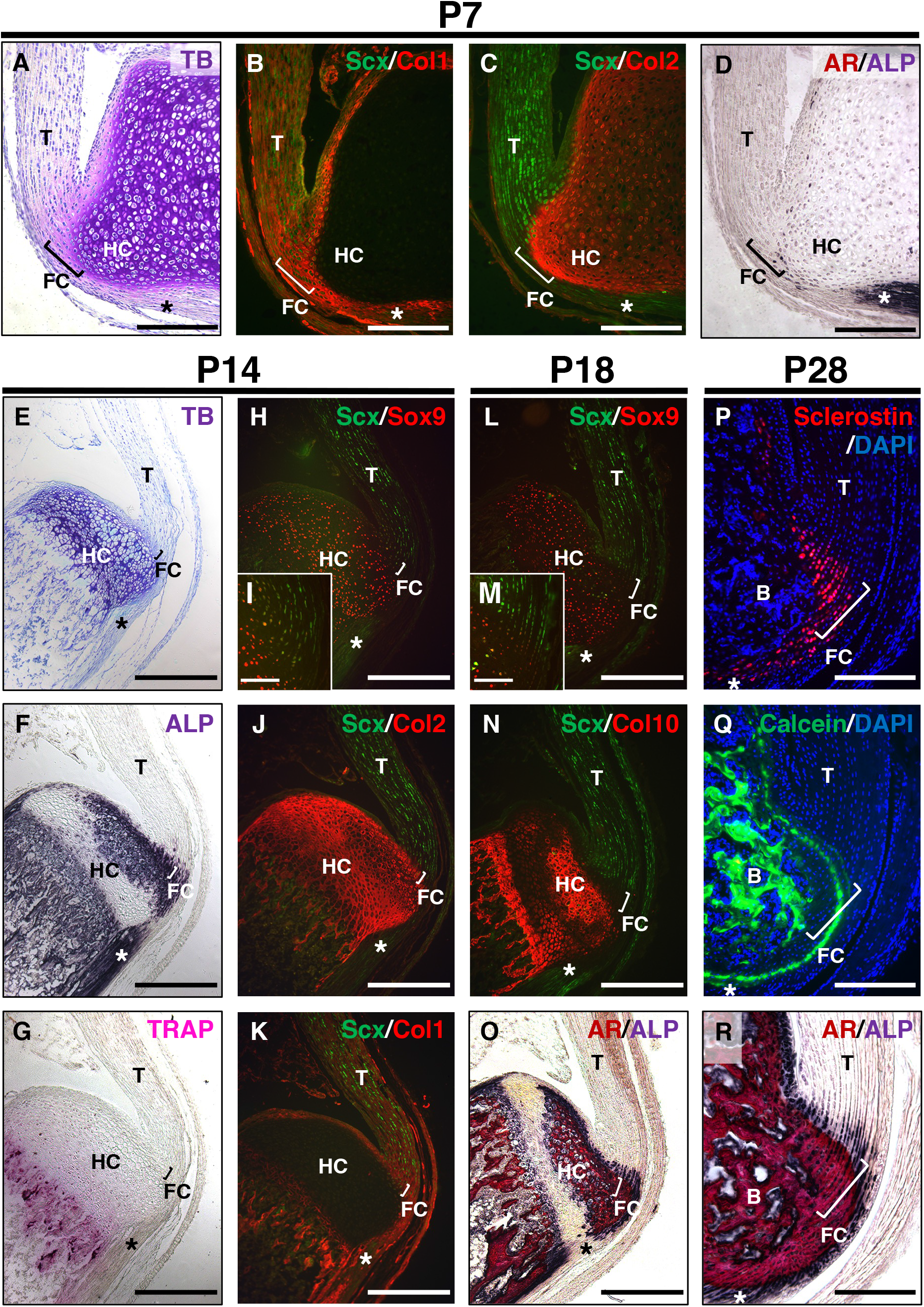
Postnatal development of entheseal fibrocartilage. Undecalcified frozen sections of the Achilles tendon enthesis were prepared from *ScxGFP* Tg mice at P7 (A-D), P14 (E-K), or P18 (L-O). Sagittal sections were processed for staining of TB (A,E), ALP (D,F,O), TRAP (G), AR (O), or immunostaining with antibodies against GFP for Scx expression (green) (B,C,H-N) and Sox9 (red) (H,I,L,M); Col2 (red) (C,J); Col1 (red) (B,K); or Col10 (red) (N). The insets (I,M) show high-magnification images of fibrocartilage immunostained with antibodies against GFP (green) and Sox9 (red). Undecalcified frozen sections of the Achilles tendon enthesis at P28 were prepared from wild-type mice administrated with Calcein at P21 and P27, respectively. Immunostaining with sclerostin (red) is shown in (P) and mineral apposition was indicated by Calcein labelling (green) in (Q). The nuclei were stained with DAPI (P,Q). AR and ALP (AR/ALP) staining are shown in (R). Square brackets indicate fibrocartilage of the enthesis. Asterisks indicate the plantaris tendon. Abbreviations: T, tendon; FC, fibrocartilage; HC, hyaline cartilage; B, bone. Scale bars: 200 µm (A-D,I,M,P-R), 400 µm (E-H,J-L,N,O).

We then examined the expression profile of sclerostin during fibrocartilaginous enthesis formation (Fig. 2). At P14, most fibrochondrocytes were present in the mineralized region adjacent to the mineralized hyaline cartilage consisting of hypertrophic chondrocytes (Fig. 2A,B). ALP^+^ cells were observed in both the unmineralized and mineralized regions (Fig. 2B). Osteocalcin (Ocn) and sclerostin were not detected in either fibrocartilage or hyaline cartilage at P14 (Fig. 2C). At P22, the secondary ossification center appeared in the epiphysis, where the mineralized hyaline cartilage was invaded by blood vessels and gradually replaced by bone (Fig. 2D,E). More intense staining of ALP was observed in the unmineralized and mineralized fibrocartilage, hyaline cartilage, and subchondral bone (Fig. 2E). Sclerostin^+^ cells were observed in the Ocn-expressing mineralized fibrocartilage at P22 (Fig. 2F). Note that cell size of fibrochondrocytes in the mineralized fibrocartilage was much smaller than that of hypertrophic chondrocytes in mineralized hyaline cartilage (Fig. 2B,E). By P45, the epiphyseal mineralized hyaline cartilage was replaced by bone, and the entheseal mineralized fibrocartilage expanded further (Fig. 2G,H). More sclerostin-expressing cells were observed in the Ocn-deposited mineralized fibrocartilage (Fig. 2I). At P84, expansion of the unmineralized fibrocartilage above the mineralized fibrocartilage was more evident in association with a decrease in ALP^+^ cells (Fig. 2J,K), but the plantaris tendon was still ALP^+^ (Fig. 2K). Sclerostin expression was largely overlapped with Ocn expression in fibrochondrocytes (Fig. 2L). In the epiphysis, hyaline cartilage was completely replaced by bone (Fig. 2H,K) and rapid mineralization of fibrocartilage without significant cellular hypertrophy are followed by expansion of the unmineralized fibrocartilage (Fig. 2J,K). In fibrocartilage, expansion of the mineralized fibrocartilage is guided by ALP^+^ cells at the mineralization front. This is followed by expansion of the unmineralized fibrocartilage after decrease of ALP^+^ cells. Sclerostin is an excellent marker for mature fibrochondrocytes located at the mineralized fibrocartilage adjacent to subchondral bone.

**Fig. 2.**
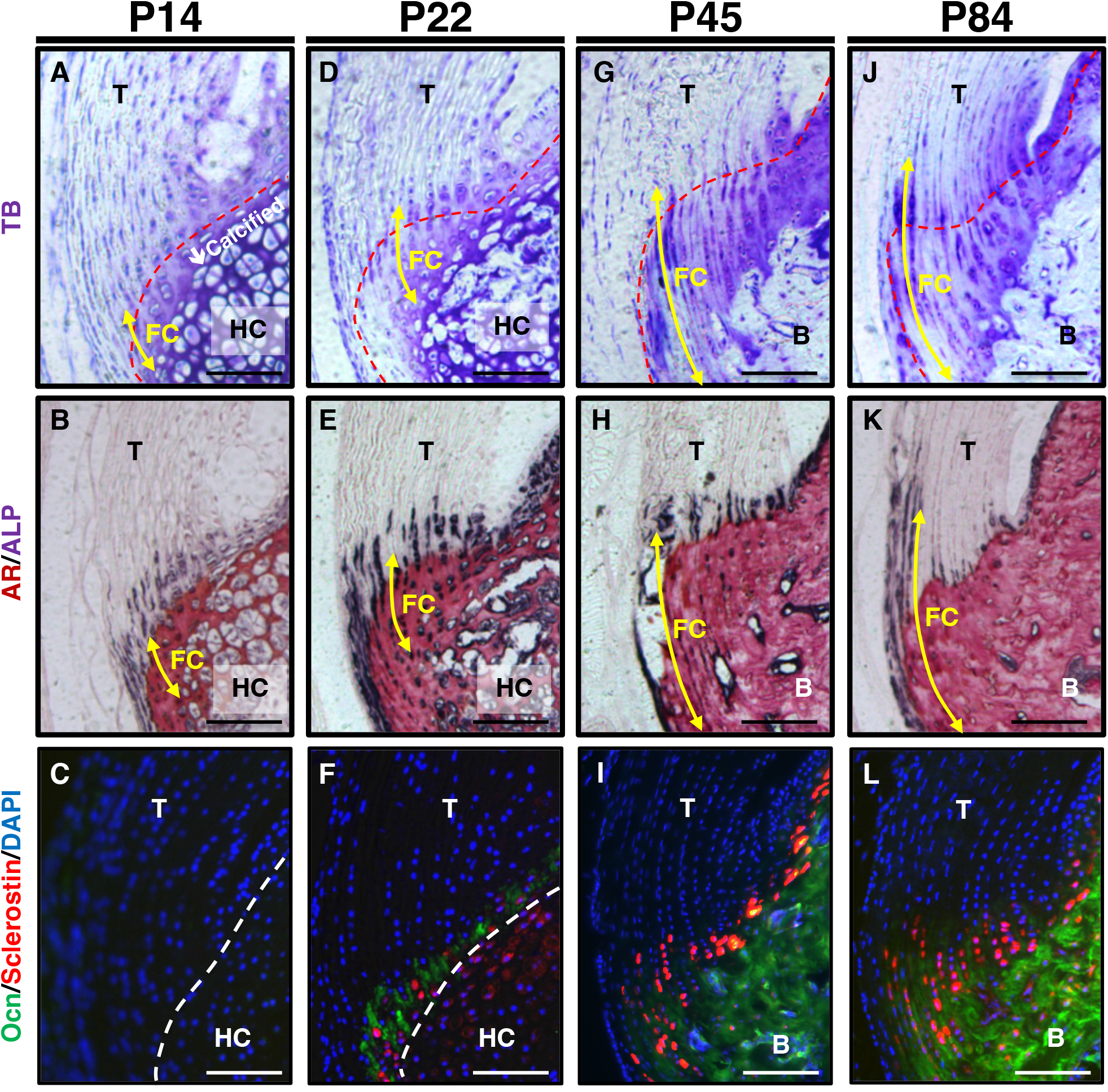
Expression of sclerostin in mature fibrochondrocytes of the Achilles tendon enthesis. (**A-L**) Undecalcified frozen sections of the Achilles tendon enthesis were prepared from *ScxGFP* Tg mice at P14 (A,B), P22 (D,E), P45 (G,H), or P84 (J,K) and wild-type mice at P14 (C), P22 (F), P45 (I), or P84 (L). Sagittal sections were stained with TB (A,D,G,J), AR/ALP (B,E,H,K) and were processed for immunostaining of Ocn (green) and Sclerostin (red) (C,F,I,L). The nuclei were stained with DAPI (blue). Yellow arrows indicate fibrocartilage. Red dashed line in TB staining indicates the tidemark between the unmineralized and mineralized fibrocartilage. White dashed line in immunostaining indicates the boundary between fibrocartilage and hyaline cartilage. Abbreviations: T, tendon; FC, fibrocartilage; HC, hyaline cartilage; B, bone. Scale bars: 100 µm.

### Defective fibrocartilage formation in association with decrease of sclerostin expression in the Achilles tendon enthesis of *Scx* deficient mice

The Achilles tendon of *Scx* deficient mice is defective and negative for Tnmd, a marker for mature tenocytes (Shukunami et al., 2006; Yoshimoto et al., 2017; Shukunami et al., 2018). Loss of *Scx* led to defective tendon and enthesis formation, which translated into impaired mechanical outcomes (Killian and Thomopoulos, 2016; Yoshimoto et al., 2017). We analyzed changes of sclerostin expression together with cartilage and blood vessel markers in the Achilles tendon enthesis of *Scx^Δ11/Δ11^* mice (Shukunami et al., 2018). At P14, in *Scx^+/+^* mice, columnar fibrochondrocytes of the enthesis were small, while chondrocytes in epiphyseal hyaline cartilage of the calcaneus became hypertrophic (Fig. 3A). Metachromatic staining with TB was observed weakly in fibrocartilage and strongly in hyaline cartilage (Fig. 3A,B). In association with defective formation of the Achilles tendon of *Scx^Δ11/Δ11^* mice, both fibrocartilaginous enthesis formation and maturation of epiphyseal hyaline cartilage were defective (Fig. 3A,B). At P28, the Achilles tendon enthesis was convex and mineralized in *Scx^+/+^* mice but rounded and unmineralized in *Scx^Δ11/Δ11^* mice (Fig. 3C-H). Fibrochondrocytes in *Scx^+/+^* mice were arranged in column along the collagen fibers connecting to the Achilles tendon (Fig. 3C), but the layer of fibrocartilage with irregularly aligned fibrochondrocytes was thin and unmineralized in *Scx^Δ11/Δ11^* mice (Fig. 3D). In *Scx^+/+^* mice, replacement of the mineralized fibrocartilage by bone was observed in the secondary ossification center (Fig. 3C,E,G). In contrast, in *Scx^Δ11/Δ11^* mice, cellular hypertrophy and mineralization of epiphyseal hyaline cartilage were delayed and vascular invasion did not occur (Fig. 3D,F,H). At P102, enthesis and epiphyseal bone formation was completed in *Scx^Δ11/+^* mice (Fig. 3I), while immature epiphysis was covered with thin ALP^+^ cells in *Scx^Δ11/Δ11^* mice (Fig. 3J).

**Fig. 3.**
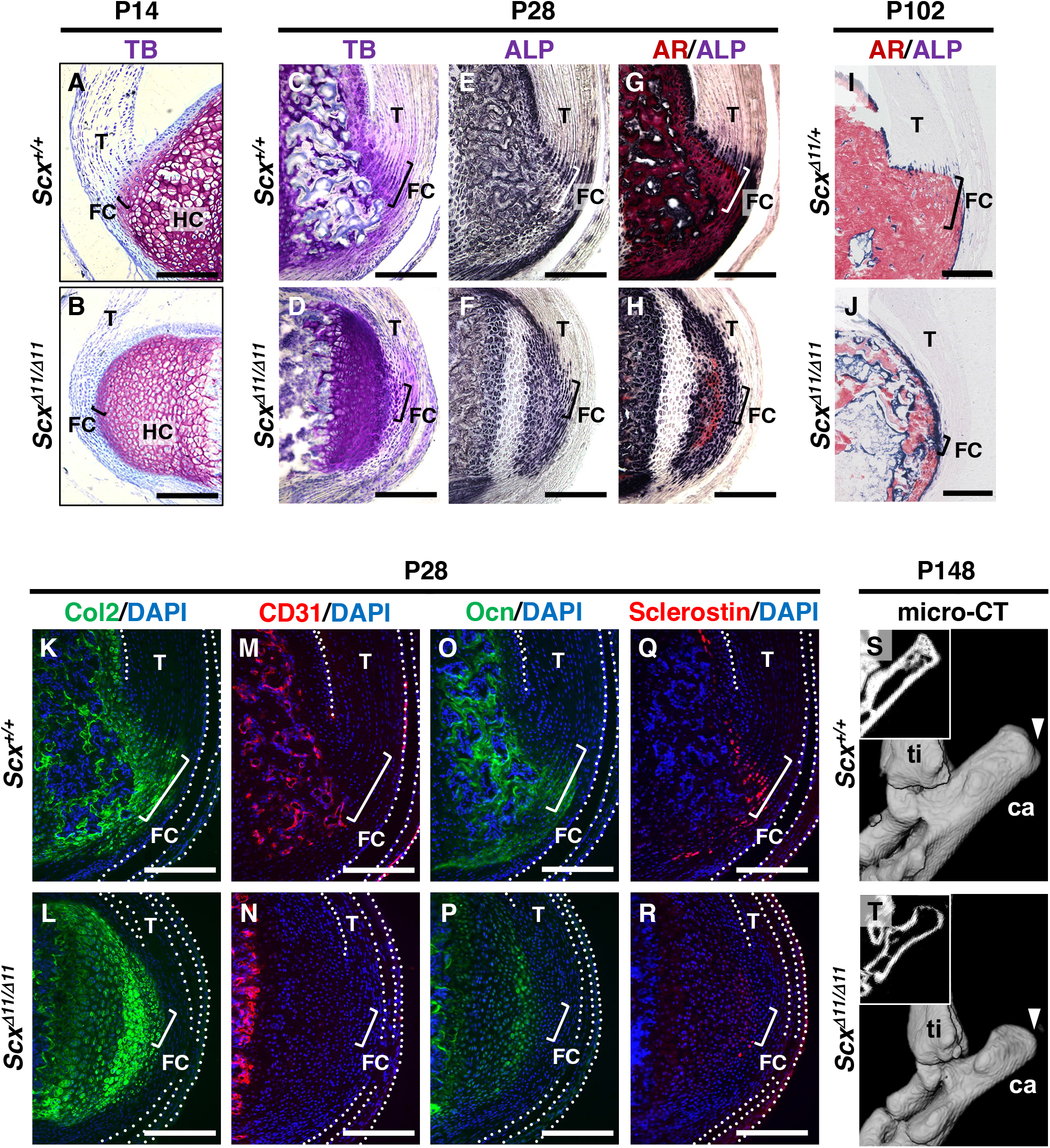
Defective fibrocartilage formation in *Scx* deficient mice. Undecalcified frozen sections of the Achilles tendon enthesis were prepared from *Scx^+/+^* (A) and *Scx^Δ11/Δ11^* (B) mice at P14 (A,B), or age-matched *Scx^+/+^* (C,E,G,K,M,O,Q) and *Scx^Δ11/Δ11^* (D,F,H,L,N,P,R) mice at P28, or *Scx^Δ11/+^*(I) and *Scx^Δ11/Δ11^* (J) mice at P102, or *Scx^+/+^* (S) and *Scx^Δ11/Δ11^* (T) mice at P148. Sagittal sections were stained with TB (A-D), ALP (E,F)/AR (G,H) or immunostained with antibodies against Col2 (green) (K,L), CD31 (red) (M,N), Ocn (green) (O,P), or Sclerostin (red) (Q,R). The nuclei were stained with DAPI (blue) (K-R). The Achilles and superficial digital flexor tendons were enclosed by white dotted lines in (K-R). Square brackets indicate the fibrocartilaginous enthesis. Micro-CT images of *Scx^+/+^* (S) or *Scx^Δ11/Δ11^* (T) mice are shown. The insets (S,T) show single planes from micro-CT images. White arrow heads in (S,T) indicate the Achilles tendon enthesis. Abbreviations: T, tendon; FC, fibrocartilage; HC, hyaline cartilage; ti; tibia; ca; calcaneus. Scale bars: 200 µm (A-R).

We then analyzed Sclerostin, Ocn, CD31 (a marker of vascular endothelial cells), Col2 localization in *Scx^+/+^* or *Scx^Δ11/Δ11^* mice at P28 (Fig. 3K-R). Wild-type sclerostin^+^ fibrochondrocytes were co-expressed with Ocn in the mineralized enthesis negative for CD31 (Fig. 3M,O,Q) but positive for Col2 (Fig. 3K). In *Scx^Δ11/Δ11^* mice, vascular invasion did not occur in epiphyseal cartilage, and sclerostin was faintly co-expressed with Ocn and Col2 (Fig. 3L,N,P,R).

Micro-CT imaging at P148 revealed that the enthesis of *Scx^Δ11/Δ11^* mice was round and undermineralized, compared with *Scx^+/+^* mice (Fig. 3S,T). These results suggest that mechanical stimulation is essential for the proper development of the fibrocartilaginous enthesis, and that sclerostin expression in the avascular mineralized fibrocartilage is closely associated with fibrochondrocyte maturation.

### Increased bone mineral density and higher stiffness in the fibrocartilaginous enthesis of *Sost****^Δ^****^26/^****^Δ^****^26^* mice

To reveal the *in vivo* function of sclerostin in the fibrocartilaginous enthesis, we analyzed *Sost* deficient mice generated by using Platinum TALENs (Sakuma et al., 2013)(Fig. S1). At P30, replacement of mineralized hyaline cartilage by bone was accelerated in *Sost^Δ26/Δ26^* mice than *Sost^Δ26/+^* mice (Fig. 4A-D). To analyze the direction and mineral apposition rate during enthesis formation, we injected two different fluorescent mineralization labels to identify the mineralization front at different time points. Calcein and Alizarin complexone were injected into *Sost^Δ26/+^* mice or *Sost^Δ26/Δ26^* at P17 and P35, respectively, and then sacrificed mice at P37 (Fig. 4E). Mineral apposition occurred from the bottom of the enthesis towards the tendon midsubstance and mineral apposition rate of *Sost^Δ26/Δ26^* mice was comparable to that of *Sost^Δ26/+^* mice (Fig. 4F,G). Sclerostin was localized to fibrocartilage and bone of *Sost^+/+^* mice (Fig. 4H) but not in those of *Sost^Δ26/Δ26^* mice (Fig. 4I) at P90. At P120, the overall number of ALP^+^ cells was decreased, but more ALP^+^ cells were observed in both fibrocartilage and bone of *Sost^Δ26/Δ26^* mice than *Sost^+/+^* mice (Fig. 4J-M). Loss of sclerostin expression in *Sost^Δ26/Δ26^* mice at P120 was also confirmed by western blotting (Fig. 4N). As shown in micro-CT images of mice at P120, the structure of the fibrocartilaginous enthesis of *Sost^Δ26/Δ26^* mice was not comparable to that of *Sost^+/+^* (Fig. 4O,P), but mineralization was enhanced in *Sost^Δ26/Δ26^* mice (Fig. 4R) than in *Sost^+/+^* mice (Fig. 4Q). Increased bone mineral density was also observed in *Sost^Δ2/Δ2^* mice compared to *Sost^Δ2/+^* mice (Fig. S2).

**Fig. 4.**
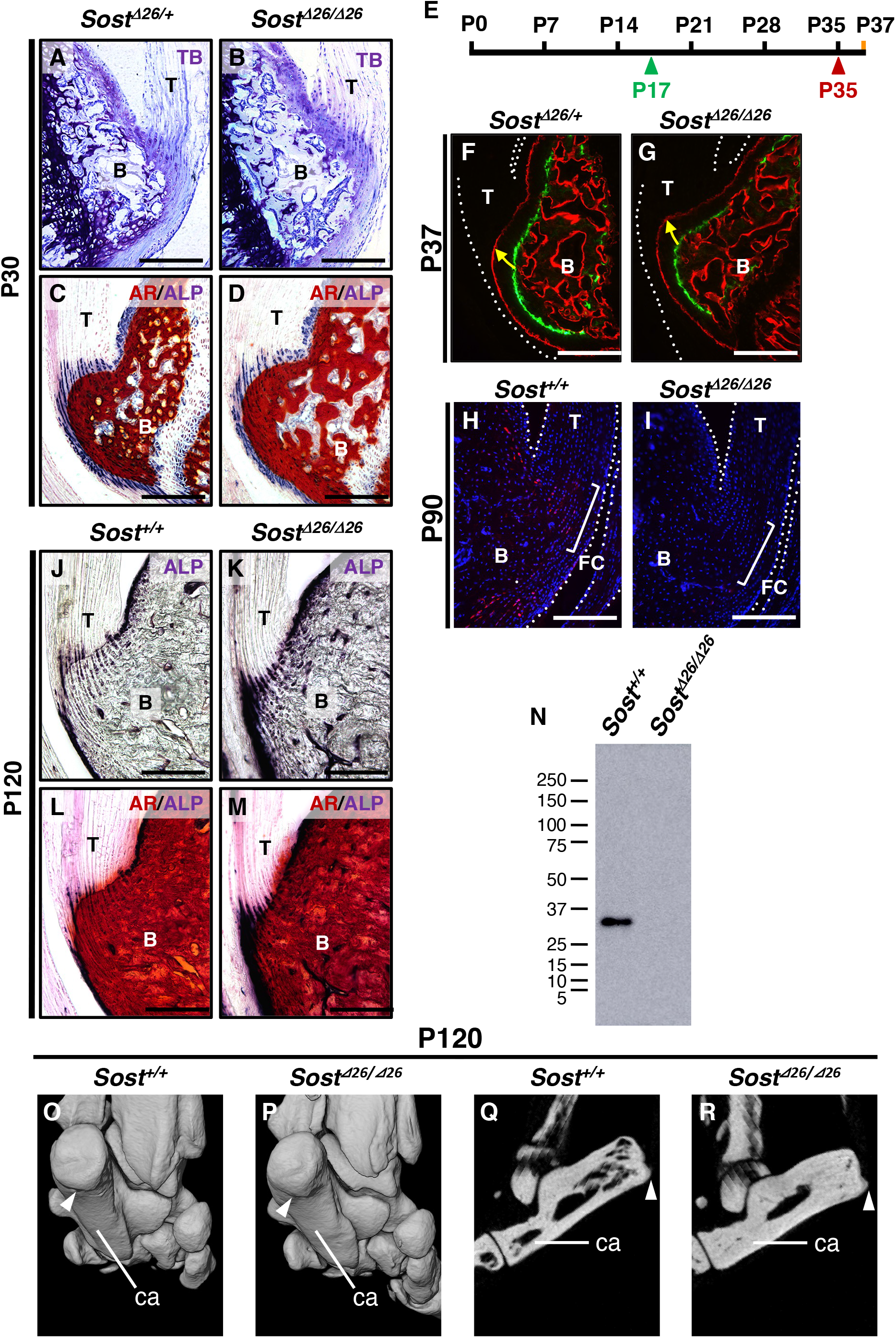
Increased mineralization of the entheseal fibrocartilage and subchondral bone in *Sost* deficient mice. Undecalcified frozen sections of the Achilles tendon enthesis were prepared from *Sost^Δ^*^26*/+*^ mice and *Sost^Δ^*^26*/*^*^Δ^*^26^ mice at P30 (A-D). Sagittal sections were stained with TB (A,B), AR/ALP (C,D). Following the experimental schedule (E), Calcein and Alizarin complexone were administrated at P17 (green arrowhead) and P35 (red arrowhead), respectively. Undecalcified frozen sagittal sections of the Achilles tendon enthesis were prepared from *Sost^Δ26/+^* at P37 (F) and *Sost^Δ26/Δ26^* mice at P37 (G). Yellow arrows denote direction of fibrocartilage mineralization and white dotted lines enclose the Achilles tendon in (F,G). Sagittal sections prepared from *Sost^+/+^* and *Sost^Δ26/Δ26^* mice at P90 were processed for immunostaining with antibody against Sclerostin (red). The nuclei were stained with DAPI (bule). Achilles and superficial digital flexor tendons were enclosed by white dotted lines in (H,I). Square brackets indicate fibrocartilage of the enthesis. Undecalcified frozen sections of the Achilles tendon enthesis were prepared from age-matched *Sost*^+*/+*^ mice and *Sost^Δ^*^26*/*^*^Δ^*^26^ mice at P120 (J-M). Sagittal sections were stained with ALP (J,K) or AR/ALP (L,M). (**N**) Western blotting was carried out for detection of sclerostin with a molecular weight of ∼28 kDa in the extract from tibia of *Sost^+/+^* and *Sost^Δ26/Δ26^* mice at P120. Micro-CT images of the left heel of age-matched *Sost^+/+^* (O,Q) and *Sost^Δ26/Δ26^* mice (P,R) at P120. Three-dimensional views of the left heel are shown in (O,P) and the sagittal plane images are shown in (Q,R), respectively. White arrowheads in (O-R) indicate the attachment site of the Achilles tendon to calcaneus bone. Abbreviations: T, tendon; FC, fibrocartilage; B, bone; ca, calcaneus.Scale bars: 200 µm (A-D,F-M).

For quantitative assessment, we scanned the calcaneus of *Sost^+/+^* mice (Fig. 5A) and *Sost^Δ26/Δ26^* mice (Fig. 5B) at P120, and performed segmentation of cortex from trabeculae in the calcaneus based on the structure and intensity of the image (Fig. 5C,D). Cortical bone thickness (Fig. 5E) and trabecular bone volume/tissue volume (Fig. 5F) of the calcaneus were significantly increased in *Sost^Δ26/Δ26^* mice compared with *Sost^+/+^* mice. To analyze mineralization of the fibrocartilaginous enthesis, we extracted the image of epiphysis of the calcaneus which includes the subchondral bone and mineralized fibrocartilage of *Sost^+/+^* (Fig. 5G) or *Sost^Δ26/Δ26^* mice (Fig. 5H) at P120 and demonstrated that bone mineral density was significantly increased in *Sost^Δ26/Δ26^* mice than in *Sost^+/+^* mice (Fig. 5I).

**Fig. 5.**
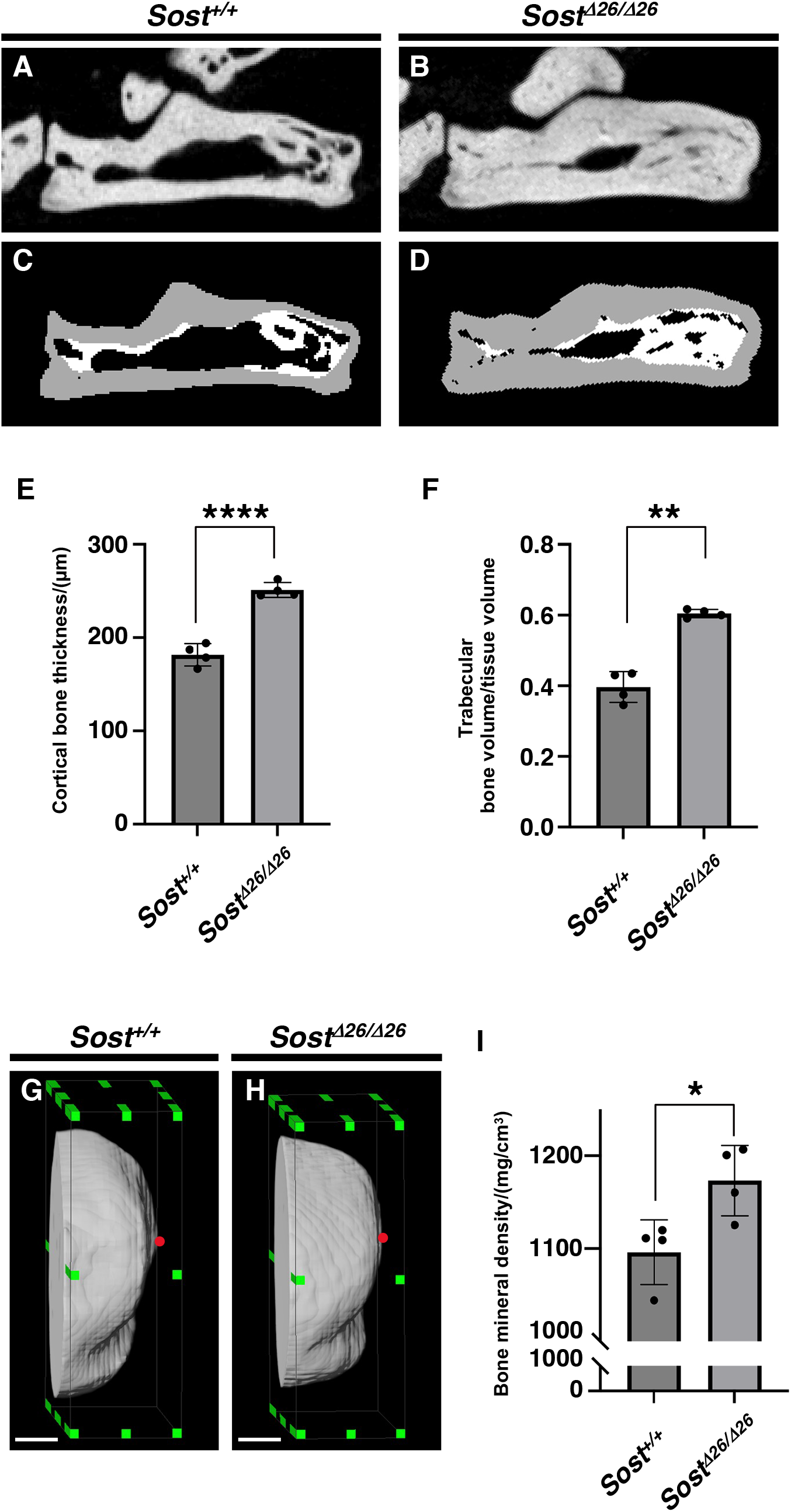
Increased bone volume and mineral density of the calcaneus of *Sost* deficient mice. The sagittal plane images of the left calcaneus of age-matched *Sost^+/+^* (A,C) and *Sost^Δ26/Δ26^* mice (B,D) at P120 are shown. The segmented areas are shown as cortical bone (gray) and trabecular bone (white), respectively (C,D). Cortical bone thickness and trabecular bone volume/tissue volume are shown in (E) and (F). Region of interest (ROI) of calcaneus for calculation of bone mineral density in age-matched *Sost^+/+^* (G) and *Sost^Δ26/Δ26^* mice at P120 (H) are indicated by green square dots. A red dot in (G,H) indicates the anterior tip of ROI. Bone mineral density in ROI is shown in (I). Scale Bars: 200 µm. *n* = 4 biological replicates per group. Data represent mean ± SEM. \**p* < 0.05 (Unpaired t test), \*\**p* < 0.01 (Unpaired t test with Welch’s t test) and \*\*\*\**p* < 0.0001 (Unpaired t test).

To analyze stiffness of the enthesis in *Sost^+/+^* and *Sost^Δ26/Δ26^* mice at P120, we conducted tissue indentation experiment using AFM (Fig. 6). Tendon, the unmineralized/mineralized fibrocartilage and bone were identified by Hoechst staining patterns under fluorescent microscope (Fig. 6A). Both the unmineralized and mineralized fibrocartilage exhibited higher stiffness in *Sost^Δ26/Δ26^* mice, with larger stiffness values, whereas tendon and bone regions did not show significant difference (Fig. 6B).

**Fig. 6.**
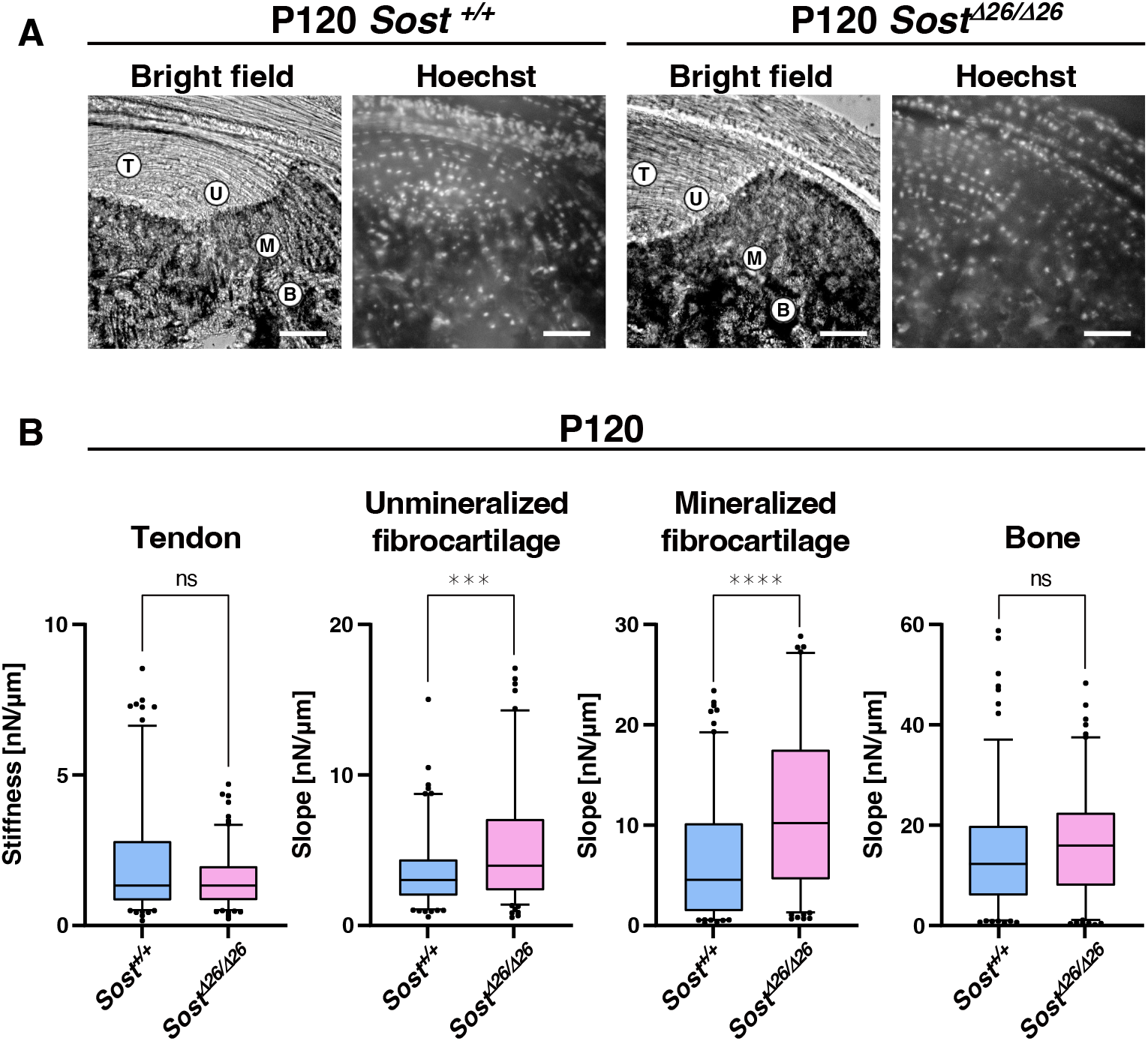
Higher stiffness of the entheseal fibrocartilage in *Sost* deficient mice. **(A)** For atomic force microscope-based tissue indentation, tendon (T), unmineralized fibrocartilage (UFC; U), mineralized fibrocartilage (MFC; M) and bone (B) regions were identified in the cryosections from P120 *Sost*^+*/*+^ and *Sost^Δ^*^26*/*^*^Δ^*^26^ mice. Scale bars: 100 µm. **(B)** Box-and-whisker plots of stiffness [nN/µm] for each tissue region. The center line, the box and the whisker indicate the median value, 25-75 percentile range and 10-90 percentile range, respectively. *n* = 50 sample points/mouse. ***p < 0.001 and ****p < 0.0001 (Mann-Whitney *U* test). *n* = 3 biological replicates per group.

These results suggested that *Sost*/sclerostin regulates the degree of mineralization and modulates the gradient of fibrocartilaginous enthesis to maintain mechanical tissue integrity.

## DISCUSSION

In this study, we demonstrated that sclerostin is an excellent functional marker for mature fibrochondrocytes of the mineralized fibrocartilage. Rapid mineralization of the unmineralized fibrocartilage is guided by ALP^+^ cells at the mineralization front, which is followed by final expansion of the unmineralized fibrocartilage after the decrease of ALP^+^ cells (Fig. 7A). In the epiphysis, fibrocartilage formation is closely associated with replacement of hyaline cartilage by bone. Mechanical forces from muscle contraction transmitted through tendon are essential for the proper development of fibrocartilaginous entheses with mature sclerostin^+^ fibrochondrocytes (Fig. 7B). Loss of function study revealed that sclerostin modulates the degree of mineralization and the stiffness profile of the fibrocartilaginous enthesis for mechanical tissue integrity. (Fig. 7C).

**Fig. 7.**
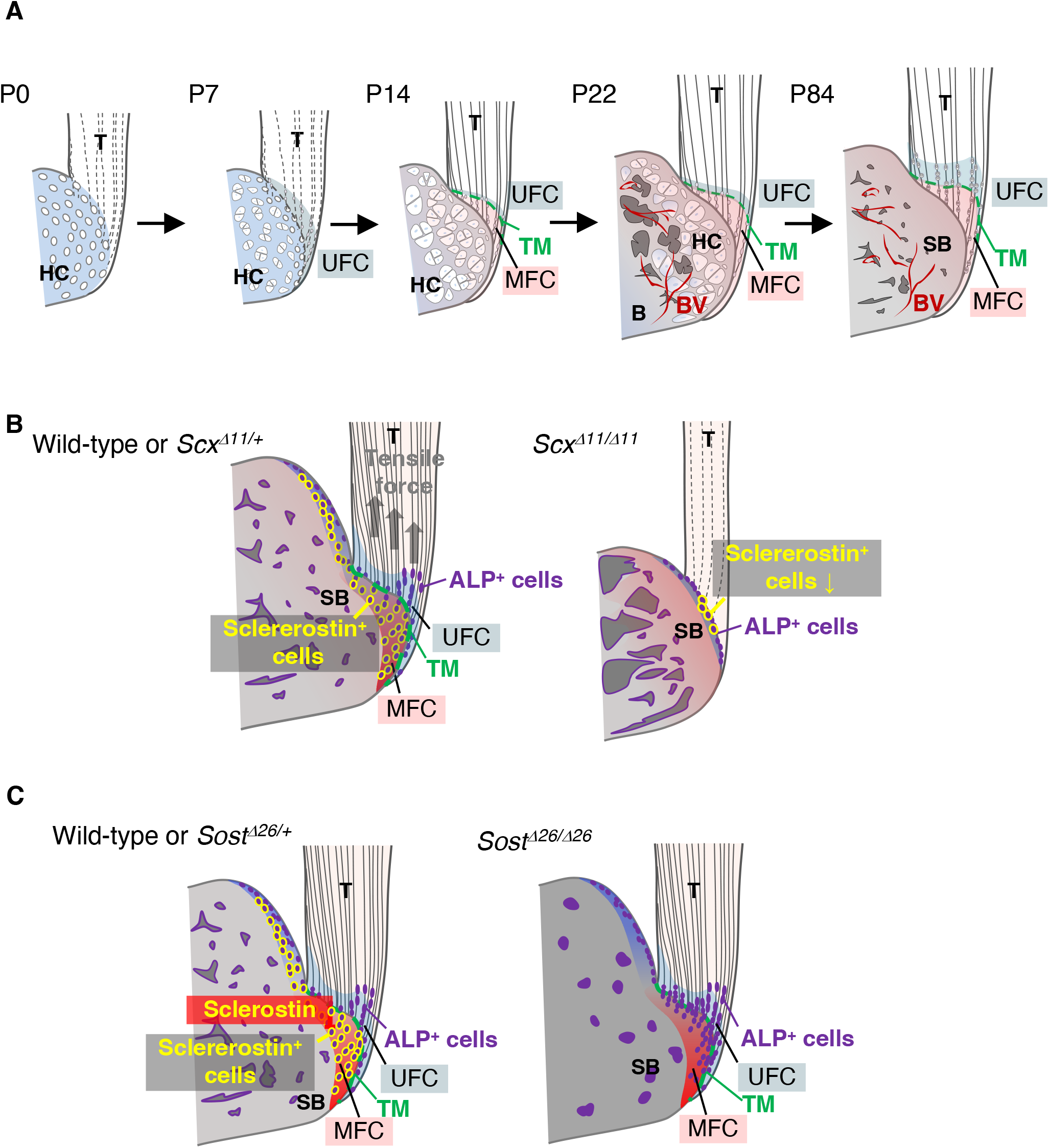
Expression and function of *Sost*/sclerostin in the fibrocartilaginous enthesis. (**A**) Schematic illustration of fibrocartilaginous enthesis formation. Development of the Achilles tendon enthesis at P0, P7, P14, P22, and P84 are presented. (**B**) Schematic illustration of fibrocartilaginous enthesis formation in wild-type or *Scx^Δ11/+^* and *Scx^Δ11/Δ11^* mice. (**C**) Schematic illustration of fibrocartilaginous enthesis formation in wild-type or *Sost^Δ26/+^* and *Sost^Δ26/2Δ6^* mice. Abbreviations: T, tendon; HC, hyaline cartilage; MFC, mineralized fibrocartilage; B, bone; BV, blood vessel; SB, subchondral bone; UFC, unmineralized fibrocartilage; TM, tidemark.

Both fibrocartilage and hyaline cartilage are generated as an avascular tissue (Benjamin and Evans, 1990; Shukunami et al., 2008; Takimoto et al., 2009), but hyaline cartilage participating in endochondral bone formation is temporary and eventually replaced by bone upon vascular invasion (Hall, 2015). In the fibrocartilaginous enthesis, mineralization rapidly progresses without a significant cellular hypertrophy. Unlike epiphyseal mineralized hyaline cartilage, mineralized fibrocartilage can be retained into adulthood due to limited resorption of the mineralized fibrocartilage via osteoclasts from the epiphyseal bone marrow (Dyment et al., 2015). Immunostaining revealed that mature mineralized fibrocartilage expressing sclerostin was devoid of CD31^+^ blood vessels. Resistance to vascular invasion from adjacent subchondral bone is thought to be an important property of the mineralized fibrocartilage that is subject to mechanical loading generated from skeletal muscle through tendon, although its molecular basis remains undetermined.

Postnatally, the primordial hyaline cartilaginous enthesis develops into the fibrocartilaginous enthesis in an orderly fashion, accompanied by cell population changes. Indeed, a recent single-cell RNA-seq study revealed that the enthesis cell populations of the shoulder rotator cuff at P11 and P18 are distinct from those at P56 (Fang et al., 2022). As shown in our present study, postnatal enthesis formation is closely associated with endochondral bone formation in the epiphysis. Hypertrophic chondrocytes within the epiphyseal calcified hyaline cartilage that is eventually replaced by bone, express a variety of growth/differentiation factors (Kronenberg, 2003) that may affect fibrocartilage formation in a paracrine manner. Indeed, indian hedgehog secreted from prehypertrophic/hypertrophic chondrocytes in epiphyseal hyaline cartilage induces Gli^+^ cells that are essential for fibrocartilage mineralization (Schwartz et al., 2015). In *Scx*-deficient mice with defective tendon and delayed maturation of epiphyseal hyaline cartilage, enthesis formation is severely impaired, probably due to a combination of reduced mechanical loading and reduced Ihh expression.

In decalcified sections, it is not possible to analyze mineralizing and mineralized chondrocytes. In this study, we used the Kawamoto’s method to keep track of the course of enthesis formation in undecalcified sections closer to the *in vivo* situation. During fibrocartilaginous enthesis formation, mineralized hyaline cartilage and fibrocartilage coexist. Of mineralizing chondrocytes in these similar but distinct cartilaginous tissue, epiphyseal hypertrophic chondrocytes are Col2^+^/Col10^+++^/Col1^-^, whereas mineralized fibrochondrocytes are Col2^++^/Col10^+^/Col1^++^. Interestingly, fibrochondrocytes, which express high levels of ALP and undergo rapid mineralization, are much smaller than hypertrophic chondrocytes in the epiphyseal hyaline cartilage. Cellular hypertrophy and Col10 expression do not appear to be directly linked to the mineralization process in enthesis formation.

It is still uncertain how the width of fibrocartilage between tendon and subchondral bone is regulated, but one important factor is undoubtedly mechanical loading. Each enthesis is uniquely mechanically loaded depending on its anatomical location, thus forming a variety of enthesis structures and sizes (Benjamin and Ralphs, 1998; Benjamin et al., 2004). In the supraspinatus tendon enthesis of *Prx1Cre^+^;Scx^flox/-^* mice, enthesis maturation is impaired and no obvious tidemark between the unmineralized and mineralized fibrocartilage is observed (Killian and Thomopoulos, 2016). Also in the Achilles tendon enthesis of *Scx^Δ11/Δ11^* mice, impaired mechanical loading led to defective fibrocartilage mineralization as well as underdevelopment of epiphyseal hyaline cartilage and subsequent bone formation, suggesting that mechanical loading is essential for proper development of these tissues.

Mineral apposition of the fibrocartilaginous enthesis and subchondral bone occur in opposite directions at the interface between the supraspinatus tendon and bone, as shown in the previous and present studies (Dyment et al., 2015). In hyaline cartilage, ALP activity is initially low in proliferating chondrocytes, but its activity increases in association with cellular hypertrophy and extracellular matrix mineralization at the primary and secondary ossification center as well as the growth plate (Cooper et al., 2013; Hallett et al., 2019). We also demonstrated that ALP^+^ cells guide mineralized fibrocartilage formation in the mineralization front that extends towards the tendon midsubstance. In the fibrocartilaginous enthesis, a unique cell population expressing Gli1, a crucial mediator of the hedgehog signaling, contributes to the postnatal development of the mineralized fibrocartilage and regeneration of fibrocartilage (Schwartz et al., 2015; Schwartz et al., 2017). Gli1^+^/ALP^+^ cells might contribute to formation of the mineralized fibrocartilage as progenitors. Similar phenomenon is observed in the secondary cartilage such as mandibular condylar fibrocartilage in which ALP^+^ progenitors rapidly differentiate into hypertrophic chondrocytes (Shibata et al., 2006). High ALP activity in these progenitor cells could allow rapid mineralization of fibrocartilage.

The mineral gradient is believed to be especially important for limiting stress concentrations and dissipating mechanical loadings at the tendon to bone interface (Schwartz et al., 2012; Tits and Ruffoni, 2021). Our study demonstrated that expansion of the mineralized fibrocartilage continued until the number of ALP^+^ cells at the mineralization front declined. It has been reported that mineralization is enhanced in chondrogenic ATDC5 cells knocked down *Sost* by lentivirus (Yamaguchi et al., 2018). Loss of *Sost* also resulted in enhanced mineralization in association with persistent expression of ALP in fibrocartilage and bone and the higher stiffness of the unmineralized and mineralized fibrocartilage as evaluated by AFM analysis. Unlike Hyp mice with enthesopathy, a murine homolog of human X-linked hypophosphatemia (Liu et al., 2018), enthesis was not expanded in *Sost* deficient mice compared with wild-type mice. Thus, sclerostin in fibrocartilage modulates the degree of mineralization and the stiffness profile to maintain mechanical tissue integrity of the enthesis without significantly affecting the morphology.

Sclerostin binds to LRP5/6 and antagonizes canonical Wnt signaling (Li et al., 2005; Yamaguchi et al., 2018), thereby promoting chondrocytes hypertrophy and subsequent ECM mineralization. Moreover, sclerostin also acts as a BMP antagonist and antagonizes BMP-6-induced ALP activity in C3H10T1/2 cells (Winkler et al., 2003). Activation of these signaling pathways has been reported in the calcaneus of *Sost* deficient mice (Krause et al., 2010). In fibrocartilage, sclerostin may negatively modulate mineral deposition of the fibrocartilage, by controlling ALP activity through suppression of Wnt and/or BMP signaling. Sclerostin regulates bone remodeling by inhibiting bone formation and promoting bone resorption (Baron and Kneissel, 2013). Since *Sost* is expressed in mature fibrochondrocytes adjacent to the subchondral bone, it may also act as a paracrine factor to participate in bone remodeling.

Entheses are hard-soft interfaces between elastic tendon and stiff hard bone with a nearly two order of magnitude stiffness mismatch and are subject to extraordinary mechanical demands (Tits and Ruffoni, 2021). Mechanosensitive properties in the tendon-enthesis-bone unit are important for mechanical tissue integrity. Scx, a functional marker for tendons and ligaments, is upregulated in response to tensile force (Takimoto et al., 2015). Sclerostin is also known to be a mechanosensitive molecule (Lin et al., 2009). Mechanical unloading leading to bone loss upregulates *Sost*/sclerostin expression in osteocytes (Lin et al., 2009), whereas mechanical loading downregulates its expression (Robling et al., 2008). Fibrocartilage is adapted to compression and/or shear stress and the deep part is compressed by the superficial part (Benjamin et al., 2006). Fibrochondrocytes embedded within mineralized matrix may sense the mechanical force to regulate their sclerostin expression. Further studies are underway how mechanical force transmitted through tendon is absorbed and converted in the fibrocartilaginous enthesis.

## MATERIALS AND METHODS

### Animals and embryos

Generation of *ScxGFP* transgenic, and *Scx^Δ11/Δ11^* strains has been previously reported (Sugimoto et al., 2013b; Shukunami et al., 2018). All animal experimental protocols were approved by the Animal Care Committee of the Institute for Life and Medical Sciences, Kyoto University, and the Committee of Animal Experimentation, Hiroshima University, and conformed to institutional guidelines for the study of vertebrates.

### Generation of TALEN-mediated *Sost* deficient mice

TALEN plasmids were constructed using the Platinum Gate TALEN Kit (Kit #1000000043, Addgene, Cambridge, MA) as previously described (Sakuma et al., 2013). To prepare TALEN mRNA, TALEN plasmids for *mSostTALEN-B-L* and *mSostTALEN-B-R* were linearized with SmaI and purified by phenol-chloroform extraction. *mSostTALEN-B-L* and *-R* mRNAs were synthesized and a polyA tail was added using the mMESSAGE mMACHINE T7 ULTRA Kit (Ambion, Austin, TX) according to the manufacturer’s instructions. After purification with the MEGAclear kit (Ambion, Austin, TX), *mSostTALEN-B-L* and *mSostTALEN-B-R* mRNAs were microinjected into the cytoplasm of fertilized eggs obtained from C57BL/6 mice. Injected eggs were transferred into the oviducts of pseudopregnant surrogate ICR female mice. Genomic DNA was extracted from the tail tips of founder mice. A 444-bp fragment of exon 1, which included recognition sites for TALENs, was amplified by PCR using primers (*Sost_GTF1*: 5′-AAGGCAACCGTATCTAGGCTGG-3′; *Sost_GTR1*: 5′-CCTCCAGGTTCTAATGCTGTGCTAG-3′). The amplified fragment was then used for direct sequencing. Sequencing was performed with the BigDye Terminator Cycle Sequencing kit and an ABI 3100 Genetic Analyzer (Applied Biosystems, Foster City, CA). For genotyping, genomic DNA was extracted from mouse ear piece and PCR was performed using a specific primer set (*Sost_GTF3*: 5’-CCCGTGCCTCATCTGCCTACTTG-3’; *Sost_GTR2*: 5’-TCTTCATCCCGTACCTTTGGC-3’). The amplified fragment was analyzed by MultiNa (SHIMADZU).

### Western blot analysis

Tibia was dissected from wild-type and *Sost^Δ26/Δ26^* female mice at P120. The isolated tissue was homogenized in RIPA buffer containing Halt Protease Inhibitor Cocktail (Thermo Fisher Scientific) and Halt Phosphatase Inhibitor Single-Use Cocktail (Thermo Fisher Scientific). The concentration of tissue extracts was quantified using a BCA Protein Assay Kit (Takara). Samples were electrophoresed on 10% TGX Stain-Free gel (Bio-Rad) and transferred to polyvinylidene fluoride membrane (Bio-Rad) using a Trans-Bio Turbo Transfer System (Bio-Rad). The membrane was incubated with anti-Mouse SOST/Sclerostin (R&D System, AF1589, 1:500) antibody in Bullet Blocking One (Nacalai Tesque) followed by horseradish peroxidase (HRP)-conjugated anti-goat IgG. Peroxidase activity was detected using SuperSignal West Pico Chemiluminescent Substrate (Thermo Fisher Scientific).

### Immunostaining

Anesthetized mice were perfused with 4% paraformaldehyde in phosphate buffered saline (PFA/PBS) containing 16.6% or 20% sucrose. The specimens were fixed in 4% PFA/PBS containing 16.6% or 20% sucrose at 4°C for 1 to 3 hours, embedded in SCEM or SCEM-L1 (Section-Lab). Undecalcified frozen sections at a thickness of 4 µm were obtained according to Kawamoto’s film method using TC-65 (Leica Microsystems) or SL-T35 (UF) (Section-Lab) tungsten carbide blades, and Cryofilm type 2C (9) or Cryofilm type 3 (16UF)(Section-Lab)(Kawamoto, 2003; Kawamoto and Kawamoto, 2021). After washing with ethanol and PBS, the sections were fixed in 4% PFA/PBS for 5 min and/or decalcified with 0.25 M ethylenediaminetetraacetic acid (EDTA) (pH 8.0). For detection of GFP, Sox9, Sclerostin, CD31, Ocn, Col1, and Col2 (for P14), sections were treated with hyaluronidase (Sigma-Aldrich) at 37°C. Sections for detection of Col2 (for P28) were treated with 1 µg/ml of protein kinase K. These sections were then fixed with 4% PFA/PBS. Sections treated with hyaluronidase were boiled in 10 mM sodium citrate buffer (pH 6.0). For detection of GFP, Sox9, Col10, sclerostin, Ocn, CD31, sections were permeabilized in 0.2% Triton X-100 in PBS. The sections were incubated with primary antibodies for 16 h or overnight, washed and then incubated with appropriate secondary antibodies conjugated with Alexa Fluor 488 or 594 (Life Technologies, Cell Signaling Technology). Nuclei were counterstained with 4 ′,6-diamidino-2-phenylindole (DAPI) (Sigma-Aldrich). The primary antibodies used were anti-GFP (Nacalai Tesque, GF090R; 1:1000), anti-Sox9 (MILLIPORE, AB5535; 1:800), anti-Col1 (ROCKLAND, 600-401-103-0.1; 1:500), anti-Col2 (ROCKLAND, 600-401-104-0.1; 1:500), anti-Col10 (abcam, ab260040; 1:250), anti-Mouse SOST/Sclerostin (R&D Systems, AF1589; 1:500), anti-Ocn (Takara, M173; 1:800), anti-CD31 (BD, 553370; 1:2000). The images were captured under a Leica DMRXA microscope equipped with a Leica DFC310 FX camera (Leica Microsystems).

### *in vivo* labeling of bone with fluorochromes

Intraperitoneal injection of Calcein (10 µg/g body weight) (DOJINDO,348-00434) and Alizarin Complexone (30 µg/g body weight) (TOKYO CHEMICAL INDUSTRY CO., LTD, A3227) diluted in 2% NaHCO_3_ were delivered to mice based on experimental designs. Labelled mice were anesthetized and perfused with 4% PFA/PBS containing 16.6% sucrose and fixed in 4% PFA/PBS containing 16.6% sucrose at 4°C for 2 hours. Undecalcified frozen sections at a thickness of 4 µm were obtained according to Kawamoto’s film method (Kawamoto, 2003; Kawamoto and Kawamoto, 2021). Nuclei were counterstained with 4’, 6-diamidino-2-phenylindole (DAPI) (Sigma-Aldrich) and the images were captured under a Leica DMRXA microscope equipped with a Leica DFC310 FX camera (Leica Microsystems).

### Histological staining

For undecalcified frozen sections, anesthetized mice were perfused with 4% PFA/PBS containing 16.6% or 20% sucrose and fixed in 4% PFA/PBS containing 16.6% or 20% sucrose at 4°C for 2 or 3 hours. Undecalcified frozen sections were stained with 0.05% TB solution at pH 4.1 (MUTO PURE CHEMICALS CO., LTD) for 5 min or TRAP staining with a TRAP/ALP Stain Kit (FUJIFILM Wako Pure Chemical Corporation) for 30 min or ALP staining with NBT/BICP solution (Roche; 1:100) for 15 min followed by AR staining prepared from 1% AR Solution at pH 6.3-6.4 (MUTO PURE CHEMICALS CO., LTD) for 5min.

### Atomic force microscope-based tissue indentation

Anesthetized male mice were perfused with PBS, and then the Achilles tendon entheses were dissected. The specimens were prepared from wild-type or *Sost^Δ26/Δ26^* mice perfused with PBS and embedded in SCEM (Section-Lab), and frozen in Hexane (Sigma-Aldrich) cooled with dry ice. Undecalcified frozen sections at a thickness of 20 µm were obtained according to Kawamoto’s film method (Kawamoto, 2003; Kawamoto and Kawamoto, 2021). Cell nuclei were stained with Hoechst 33342. For AFM-based tissue indentation (H-J Butt, 1995; Ryo Ichijo, 2022), JPK BioAFM NanoWizard 3 (Bruker Nano GmbH) was employed. The AFM system was mounted on a bright field/fluorescence microscope (IX81; Evident Co.). AFM cantilevers (TL-CONT; spring constant 0.2 N/m; Nanoworld AG) were modified with glass beads with a diameter of 10 μm and calibrated using the thermal noise method (H-J Butt, 1995). To identify the tendon, fibrocartilage and bone regions, Hoechst-stained tissue sections were observed by IX81 microscope. Especially, to classify the unmineralized and mineralized fibrocartilage regions, AR-stained sections from the same tissue were referred. For AFM-based indentation, the piezo displacement speed and the sampling rate were set as 3 μm/s and 4,000 Hz, respectively. The obtained indentation force (*F*) versus depth (*h*) curve was smoothed using a moving average for 10 datum points before and after each averaging point. Stiffness [nN/μm] were estimated by linear regression for the (*F*, *h*) datum points within an indentation depth range of 0 nm ≤ *h* ≤ 50 nm.

### Skeletal imaging by micro-computed tomography analysis

Mice at P58 and P148 were fixed with 4% PFA/PBS or 95% Ethanol and subjected to scans with inspeXio SMX-90CT (SHIMADZU). Anesthetized mice at P120 were perfused with 4% PFA/PBS or PBS and soaked in 99.5% ethanol (Wako, 057-00456). Mice were analyzed by inspeXio SMX-90CT Plus (SHIMADZU) in 99.5% ethanol (Wako, 057-00456) with a 90 kV source voltage and 110 µA source current and a resolution of 0.026 mm/pixel (*n*=4 biological replicates for each group). Two-and three-dimensional reconstruction was performed using Amira 3D software version 2021.1 (Thermo Fisher Scientific). Bone mineral density was calculated from the equation derived from least-squares methods with 5 plots using hydroxyapatite phantom with 100 mg/cm^3^, 200 mg/cm^3^, 300 mg/cm^3^, 400 mg/cm^3^, 500 mg/cm^3^ (RATOC, No06-U5D1mmH) was separately scanned on the same day under the same conditions for the samples.

### Statistics

All statistical analyses were carried out using Microsoft Excel or GraphPad Prism version 9 (GraphPad Software, LLC). Data are presented as mean ± SEM. Comparisons were performed using unpaired t test (for cortical bone thickness and bone mineral density) or unpaired t test with Welch’s correction (for trabecular bone volume/tissue volume) or Mann-Whitney *U* test (for AFM-based tissue indentation) to determine significance between groups. The level of significance was set at *P* < 0.05.

## Acknowledgements

We thank Dr. Yoshitaka Kameo, Dr. Masaki Takechi, and Dr. Sachiko Iseki for helpful discussion.

## Competing interests

The authors declare no competing or financial interests.

## Author contribution

C.S. designed the study and supervised the project. S.Y., Y.Y., K.I., K.M., A.T., S.H., X.Y., H.W., T.S., T.Y., G.K., C.S. performed the experiments. C.S., S.Y., Y.Y., K.M., K.I., A.T., S.H., T.Y., K.U., S.M., K.T., D.D., and T.A. analyzed and interpreted the data. C.S., S.Y., Y.Y., and K.I. prepared the manuscript.

## Funding

This work was also supported by JSPS Grants-in-Aid for Scientific Research (Grant Numbers JP21H03107, JP18H02966, JP21KK0161, JP17K17092), JST-CREST, Grant Number JPMJCR22L5, Phoenix Leader Education Program for Renaissance from Radiation Disaster funded by the Program for Leading Graduate Schools, the Frontier Development Program for Genome Editing funded by the Doctoral Program for World Leading Innovative and Smart Education, JST SPRING, Grant Number JPMJSP2132, and the Cooperative Research Program of Institute for Frontier Life and Medical Sciences, Kyoto University.

## Notes

### Competing Interest Statement

The authors have declared no competing interest.

