## Supplemental Information for "Sclerostin modulates the degree of mineralization and the stiffness profile of the fibrocartilaginous enthesis for mechanical tissue integrity"

### Supplementary Figure 1.

**A**

MQPSLAPCLICLLVHAAFCAVEG**QGWQAFRNDATEVIPGLGEYPEPPPENNQTMNRAENG** 60

↳ **sclerostin**

**GRPPHHPYDAKDVSEYSCRELHYTRFLTDGPCRSAKPVTELVCSGQCGPARLLPNAIGRV** 120

**KWWRPNGPDFRCIPDRYRAQRVQLLCPGGAAPRSRKVRLVASCKCKRLTRFHNQSELKDF** 180

**GPETARPQKGRKPRPGARGAKANQAELENAY** 211

**B**

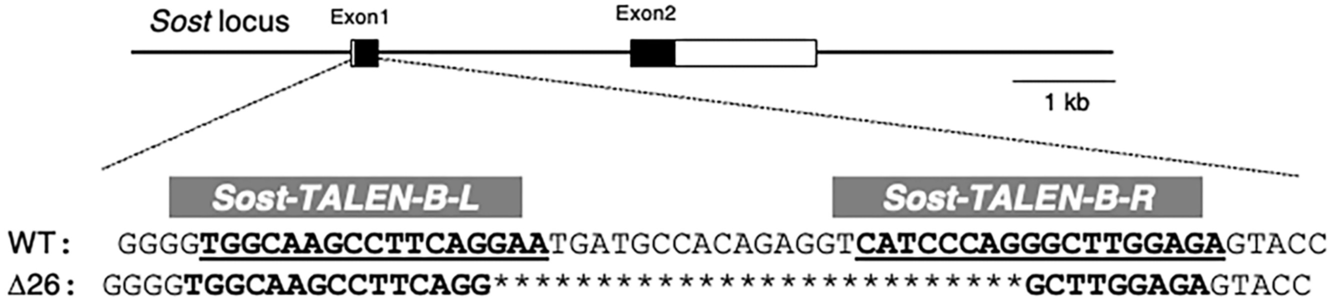


**C**

WT

**ATG** CAG CCC TCA CTA GCC CCG TGC CTC ATC TGC CTA CTT GTG CAC GCT GCC
**M** Q P S L A P C L I C L L V H A A 17
TTC TGT GCT GTG GAG GGC CAG GGG TGG CAA GCC TTC AGG AAT GAT GCC ACA
F C A V E G Q G W Q A F R N D A T 34
GAG GTC ATC CCA GGG CTT GGA GAG TAC CCC GAG CCT CCT CCT GAG AAC AAC
E V I P G L G E Y P E P P P E N N 51

Δ26

**ATG** CAG CCC TCA CTA GCC CCG TGC CTC ATC TGC CTA CTT GTG CAC GCT GCC
**M** Q P S L A P C L I C L L V H A A 17
TTC TGT GCT GTG GAG GGC CAG GGG TGG CAA GCC TTC AGG GCT TGG AGA GTA
F C A V E G Q G W Q A F R A W R V 34
CCC CGA GCC TCC TCC **TGA**
P R A S S 39

**D**

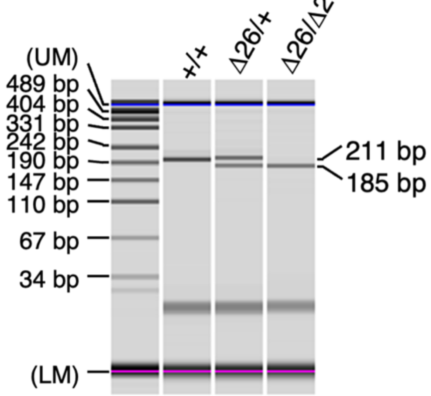

**Fig. S1. Amino acid sequences of mouse sclerostin and frameshift mutations with early stop codon created by TALENs.**

(**A**) Mouse sclerostin consisting of 211 amino acids is shown. The putative signal peptide is enclosed. Sclerostin shown in bold is a protein containing 188 amino acids from Q24 to Y211. **(B)** The genomic structure of *Sost* and TALEN target sequences in the mouse *Sost* locus. The left (L) and right (R) binding regions of *mSost-TALEN-B* are indicated by bold and underlined text. Sequences of wild-type (WT) and a founder mouse generated by microinjection of *mSost-TALEN-B-R/L* mRNAs. Nucleotide deletions are indicated by asterisks. (**C**) Sequence of a mouse line with a 26-base pair deletion. The initiation codon and the premature stop codon are highlighted in red and black. **(D)** Genotyping PCR was performed using DNA of ear piece with primers as described in Materials and Methods. The targeted *Sost*$^{\Delta 26}$ allele (185 bp) and wild-type allele (211 bp) are distinguished.

### Supplementary Figure 2.

A

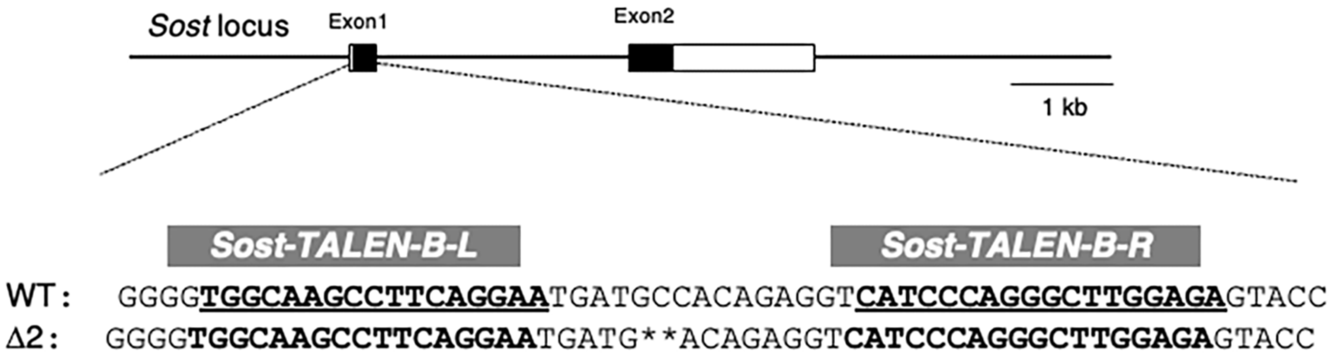


B

Δ2

ATGCAGCCCTCACTAGCCCCGTGCCTCATCTGCCTACTTGTGCACGCTGCCTTCTGTG
CTGTGGAGGGCCAGGGGTGGCAAGCCTTCAGGAATGATG**ACAGAGGTCATCCCAGG
GCTTGGAGAGTACCCCGAGCCTCCTCCTGAGAACAACCAGACCATGAACCGGGCGGAG
AATGGAGGCAGACCTCCCCACCATCCCTATGACGCCAAAG

C

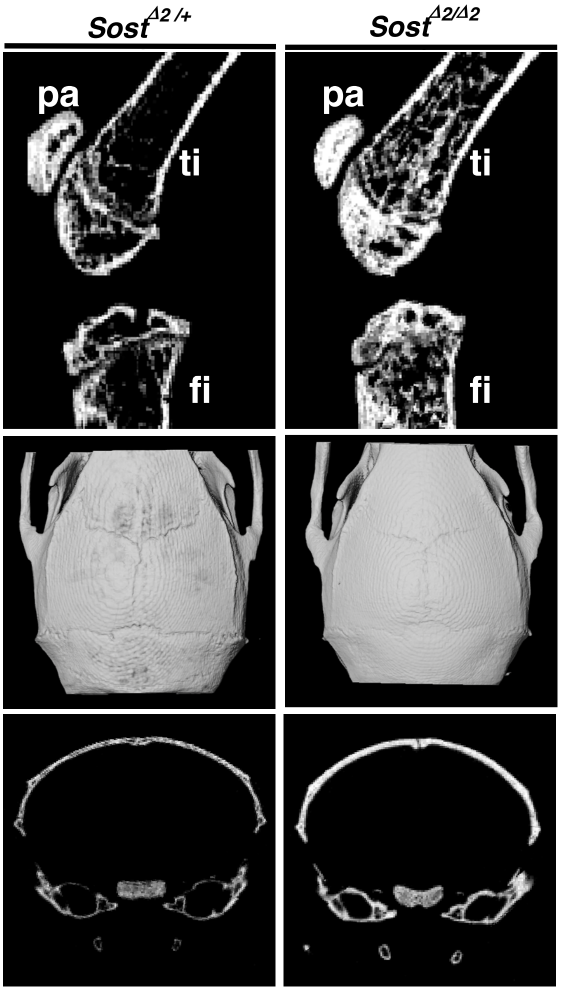

**Fig. S2. Increased bone mass in *Sost*$^{\Delta 2/\Delta 2}$ mice.**

**(A)** The genomic structure of *Sost* and TALEN target sequences in the mouse *Sost* locus. The left (L) and right (R) binding regions of *mSost-TALEN-B* are indicated by bold and underlined text. Sequences of wild-type (WT) and a founder mouse generated by microinjection of *mSost-TALEN-B-R/L* mRNAs. Nucleotide deletions are indicated by asterisks. (**B**) Sequence of a mouse line with a 2-base pair deletion. The initiation codon and the premature stop codon are highlighted in red and black. **(C)** Micro-CT images of knee joint and calvaria at P58 *Sost*$^{\Delta 2/+}$ mice and *Sost*$^{\Delta 2/\Delta 2}$ mice are shown. Abbreviations: pa, patella; ti, tibia; fi, fibula.
